# Mapping Cumulative Impacts to Coastal Ecosystem Services in British Columbia

**DOI:** 10.1101/698365

**Authors:** Gerald G. Singh, Ian M.S. Eddy, Benjamin S. Halpern, Rabin Neslo, Terre Satterfield, Kai M.A. Chan

## Abstract

Ecosystem services are impacted through restricting service supply, through limiting people from accessing services, and by affecting the quality of services. We map cumulative impacts to 8 different ecosystem services in coastal British Columbia using InVEST models, spatial data, and expert elicitation to quantify risk to each service from anthropogenic activities. We find that impact to service access and quality as well as impact to service supply results in greater severity of impact and a greater diversity of causal processes of impact than only considering impact to service supply. This suggests that limiting access to services and impacts to service quality may be important and understanding these kinds of impacts may complement our knowledge of impacts to biophysical systems that produce services. Some ecosystem services are at greater risk from climate stressors while others face greater risk from local activities. Prominent causal pathways of impact include limiting access and affecting quality. Mapping cumulative impacts to ecosystem services can yield rich insights, including highlighting areas of high impact and understanding causes of impact, and should be an essential management tool to help maintain the flow of services we benefit from.

## 1. Introduction

Humanity’s great and growing influence on the planet demands an increased understanding of how multiple activities cumulatively affect the human benefits and values associated with the environment (1, 2). In part due to their ease of comprehension and display of multiple human stressors at once, impact mapping has gained much traction in environmental science and management (3-11). However, impact mapping studies generally reflect how human activities affect species and habitats, neglecting thus far how multiple activities cumulatively affect ecosystem services – the processes by which nature renders benefits for people (12). Understanding impacts on ecosystem services would allow for a representation of multiple societal benefits from the environment, enabling targeted management on specific ecosystem services. Assessments of impacts on ecosystem services could allow us to establish baseline knowledge of the ecosystem services and geographic areas facing the greatest impact, as well as help evaluate and plan for emerging impacts from local and global stressors (such as from future oil spills and climate change, respectfully).

While any human activity that impacts ecosystems has the potential to impact ecosystem services, ecosystem services includes dynamics beyond the ecological production of potential benefits to people (13). Impacts to ecosystem services are potentially different than impacts to ecosystems. While impacts to ecosystems have (usually adverse) consequences for populations, species, and structure of ecosystems, impacts to ecosystem services ultimately negatively affect human wellbeing. For example, pollution might not affect shellfish growth, but it may lead to aquaculture tenure closure for health concerns, or might affect the taste of shellfish caught at polluted sites (14). Changes to the enjoyment of shellfish aquaculture, in this case, are not a result of changes in the biophysical supply of the service but in the change to either the access to or quality of the ecosystem service (i.e. environmental impact may not mean ecosystem service impact).

Impacts to ecosystem services can be characterized at each step in the ecosystem services ‘cascade’ (15), with human impacts potentially affecting supply (the biophysical components that produce ecosystem services), service (the ability of people to access and benefit from a service), and value (people’s preferences for ecosystem services, 13). Reframing the previous example, pollution impacts the service (the ability of people to access shellfish for food through legal restriction) or the value (the palatability of the shellfish), but the growth of shellfish (the supply) is unaffected. Conversely, sedimentation may smother and impact the population of shellfish (the supply) without affecting the quantity of shellfish sold, while people’s access (the service) to and palatability of shellfish (the value) is unchanged. In the former case what may not be considered an environmental impact (to shellfish) would be considered an ecosystem service impact. In this latter case what would be considered an environmental impact is not considered an impact to an ecosystem service. While this cascade is useful for parsing out the dynamics of impacts to ecosystem services, the relative importance of these factors (supply, service, and value) in regulating impact to ecosystem services is not known.

The ecosystem service cascade has potential repercussions for impact mapping. Because delivery of ecosystem services to people requires both the provision of services through biophysical means (the supply) and delivery to people (service) that demand those services (value), maps of ecosystem services may be more restricted in space than maps of total service supply. The few existing studies mapping cumulative impacts to ecosystem services do so using human use and landscape proxies of ecosystem services (11, 16). Recently, spatial models have been created that utilize production functions for ecosystem services, relating landscape features important for ecosystem services, as well as spatial social data on human use of the environment, to generate maps of ecosystem services on the coast (17, 18). These models provide a more precise and comprehensive mapping of ecosystem services, including those without close human use proxies.

Beyond spatial representation, the ecosystem service cascade influences the metrics we use to measure impact. Existing frameworks of impact to ecosystem services characterize change in the underlying ecosystem as the principal (and sometimes sole) driver of impact, with human beneficiaries of services largely subject to changes in ecosystem service supply (19-22). However, including metrics that consider impacts to the service and value of ecosystem services can modify our understanding and measurements of impact. Maps of species and habitat may be sufficient to approximate impacts to ecosystem services when the mechanisms of impact operate mostly through biophysical supply. However, changes to people’s access, use, and perceived quality of service may also be important for understanding impacts to ecosystem services. Frameworks of impact to ecosystem services should therefore be updated to reflect these types of impacts (23, 24). As a consequence of, including service and value dimensions, the potential causal mechanisms of impact on ecosystem services expands and ultimately leads to greater uncertainty in conceptualizing impact (25). Thus, explicitly outlining causal mechanisms of impact is an important complement to measuring impact.

Here we model human impacts to specific ecosystem services on coastal British Columbia to identify areas of high impact in consideration of the ecosystem service cascade and advance the understanding of impacts to ecosystem services. Coastal British Columbia is an area renowned for its scenery and productivity, contributing greatly to the economy, sense of place and other values important to residents and visitors (26, 27). Maps of cumulative impacts to coastal British Columbia ecosystems have been produced (8, 10, 28). This work, alternately, does so for ecosystem services themselves. We ask: 1) Which ecosystem services face the greatest cumulative impact in coastal British Columbia?; 2) What human activities pose the greatest threat to what ecosystem services?; 3) Where are ecosystem services under greatest threat?; 4) How do the answers to the first three questions change if measures of service and value are considered or left out (i.e. consider impacts to the ecosystem service cascade versus only to supply)?; 5) How may projected future impacts affect ecosystem services?; 6) What is the relative importance of metrics of service supply, service, and value to impacts on ecosystem services?; 7) What are the main causal pathways of impacts that affect the ecosystem services? Together, addressing these questions builds on established methods to map cumulative impacts using geospatial data and expert derived estimates of ecological vulnerability (4, 29).

## 2. Methods

Mapping and quantifying impacts to ecosystem services requires both (1) understanding the location and intensity of human activities co-occurring with ecosystem services (i.e., the ‘footprint’ of human activities) and (2) the risk each activity poses to each ecosystem service (19, 20). Measuring impact as a product of risk (the potential of an activity to impact an ecosystem service where they co-occur) and the co-occurrence of activities (with measured intensities) and ecosystem services follows the conceptual structure of cumulative impact mapping (33).

The analysis consisted of five main steps. First, we mapped eight ecosystem services using InVEST models and spatial data available for the region. Second, we assembled spatial data for 21 human activities and stressors that potentially impact ecosystem services. Third, we derived risk scores for each service-activity (risk of activity *x* on service *y*) combination under current conditions (within the last 10 years) via an expert elicitation process. These risk scores were calculated by expert derived estimates of risk criteria and criteria weights, then combined with data on human activities and stressors generates impact scores. Fourth, we overlaid impact scores with ecosystem service to assess the cumulative impacts of all available activities on each service. The resulting maps allowed us to answer where ecosystem services were under greatest impact, which ecosystem services were most impacted, and by what human activities or stressor. The expert scores allowed us to distinguish impacts on ecosystem service supply from impacts on service and value considerations. We compared maps of total impact with maps that only incorporated impacts on ecosystem service supply to explore the importance of service and value dimensions of ecosystem service risk. We used expert elicitation to estimate the risk of key climate change and potential oil spills to ecosystem services in the future. To further explore how ecosystem services across supply, service and value, we asked experts to detail the causal pathways of impacts to ecosystem services. We detail each step below.

### 2.1 Spatial Representation of Ecosystem Services

We mapped eight different ecosystem services using InVEST models (17, 18), or, when the models were unnecessary, using data on the extents of ecosystem services. The ecosystem services we mapped were: commercial demersal fisheries, commercial pelagic fisheries, finfish aquaculture, shellfish aquaculture, marine recreation, coastal aesthetics, coastal protection, and potential wave and tidal energy generation. We modeled “potential” energy generation because British Columbia currently does not have wave and tidal energy operations, but there is interest in harnessing this energy supply.

The InVEST tool has tiered models for mapping ecosystem services based on different levels of data availability. The highest tier InVEST models are capable of quantifying and calculating monetary values of ecosystem services within the area that people use them (17). Due to data limitations, we were prevented from modeling ecosystem services at the most refined tier, but we could produce maps of the extent of human use of ecosystem services for all eight across coastal BC. We used the base InVEST models for fisheries and recreation maps whereby overlapping maps of different activities creates the resulting service model. We modeled coastal aesthetics with InVEST by calculating the viewshed from sites of recreation and human habitation. This model considers bathymetry and topography to calculate the viewshed. We modeled coastal protection with InVEST by mapping the parts of the coast protected by vegetation, kelp, and erosion-resistant substrate (not mapped are areas of the coast without protection). We did not use InVEST to map potential renewable energy and aquaculture, as we opted to use instead the publicly available spatial data on wave and tidal energy areas of interest along the BC coast, as well as the locations of shellfish and finfish aquaculture. See the Appendix A for detailed descriptions of ecosystem service model parameterization.

### 2.2 Spatial Representation of Impacting Activities

We assembled spatial data layers for 21 different activities and stressors, including activities and stressors related to fisheries, coastal commercial industries, land based pressures, and climate change impacts (these broad categories derived from (8), see Table S2). These spatial data layers included the spatial range of each human activity and stressor, as well as the intensity of each activity within its range (for example, how many ships were using a particular shipping lane). Many human activities, such as fishing, are important ecosystem services while also contributing impacts towards ecosystem services (30). Some categories are therefore represented in both ecosystem services as well as activities and stressors that cause impact. We treat ecosystem services as broad categories (such as demersal vs pelagic fisheries) and activities and stressors that cause impact more specifically (such as demersal destructive fisheries) because of differential impacts across activities, and because experts indicated that broad types of ecosystem services (such as various benthic fisheries, or various pelagic fisheries) are impacted in similar ways. Many of the data layers of human activities and stressors that cause impact were adapted from a previous cumulative impact study by Ban et al. (8) supplemented with data from British Columbia Marine Conservation Analysis (31) and GeoBC (32). We compiled the 25 fisheries used by Ban et al. (8) into five categories (demersal destructive, demersal non-destructive, pelagic low bycatch, pelagic high bycatch, and recreational fishing) of fisheries that cause impact because the number of data layers influences the overall cumulative impact scores (8), and we did not want to overly bias impact based on fisheries scores. This dataset considers the area of influence of each human activity, with the extent of each area of influence dependent on prominent stressors associated with each activity. We also included current climate stressors adapted and updated from Halpern et al. (33) global map (see Table S3 for data sources).

### 2.3 Risk Assessment

Following the cumulative impact mapping approach first demonstrated by Halpern et al. (33), we overlay maps of impacting activities on ecosystem services, and calculate impact of activities by combining the spatial data of activity intensity with a measure of risk from a standard unit of human activity to a given ecosystem service. Calculating the quantitative estimates of risk and the model of cumulative impact were adapted from Halpern et al. (33) and are described in detail in the Supplementary Methods. Below we describe the expert elicitation process used to generate risk scores and summarize the cumulative impacts model that build from the ecosystem service maps, human activity maps, and the risk assessment.

#### 2.3.1 Expert Elicitation for Risk Scores

Our quantitative estimates of risk represent the potential impact that a unit of an activity poses to an ecosystem service when an activity co-occurs with an ecosystem service. To calculate the risk scores we relied on expert judgement, due to pervasive data gaps (see Table S4 for a description of activities and stressors for risk quantification). We adapted the mail-in and phone expert survey used in Teck et al. (29) used to quantify ecosystem vulnerability to different human activities through ranking and quantification exercises, and adapted it for ecosystem services (survey description below). We used an online survey because it allowed us to reach all experts using a common platform. The diversity of ecosystem services and the large number of risk values precluded individual surveys, workshops, and other elicitation methods (34). We invited a total of 437 experts to take part in the full survey (quantifying risk criteria, future risk criteria, generating risk criteria weights, and outlining mechanistic pathways of impacts), but 217 did not respond to the survey invitation, resulting in 220 potential expert responses (we could not determine whether these were appropriate experts who chose not to respond or if they did not receive or see the invitation). Of the resulting 220, 112 self-indicated that their level of expertise was not sufficient to quantify risk though all 220 did provide responses on the mechanisms of impact. After accounting for non-responses and self-identification, we were left with a pool of 108 confirmed potential experts. Of this pool, 44 provided quantitative results on the survey (a 40.7% response rate).

Experts were selected by reviewing the literature of the various chosen ecosystem services in British Columbia and identifying authors of relevant studies. Authors and studies were identified through ISI Web of Knowledge with a focus on recruiting experts with subject-expertise in specific (or multiple) ecosystem services specifically within BC. We allowed participants to self-organize for chosen ecosystem services (some indicating their expertise for multiple ecosystem services), and they provided responses for all ecosystem services they presumed themselves experts on in BC. Risk estimates were compiled for commercial fisheries generally (instead of demersal and pelagic commercial fisheries), and we elicited risk scores for commercial aquaculture generally (instead of shellfish and finfish aquaculture) because the fisheries and aquaculture experts indicated their expertise pertained to these ecosystem services across their subcategories.

Experts were tasked with quantifying risk according to seven criteria, building on those used in Teck et al. (29). The criteria encompassed exposure (area of influence, frequency of impact and recovery time, Table S4) and consequence (magnitude of impact on ecosystem service production, ecological extent of impact, effects to access and effects to perceived quality, Table S5). The consequence criteria include considerations across the ecosystem service cascade. Impacts to supply dimensions are represented by magnitude of impact on service production and ecological extent, impacts to service dimensions are represented by effects to access, and impacts to value dimensions are represented by effects to perceived quality. Experts were instructed to consider current risk of activities to ecosystem services (within the last 10 years). For potential energy generation, only one expert provided these quantitative measurements (though others provided other information on potential energy generation) so quantitative results for this ecosystem service should be considered tentative, and future research should be taken to verify findings here. For all other ecosystem services, there were ≥3 experts providing measurements, consistent with expert input on previous cumulative mapping studies (4, 29, 35). While we acknowledge that expert input can carry high uncertainty, expert input was the best option present give that no empirical results exist as an alternative, though empirically quantifying impacts to marine systems is a priority research area (35). Despite this limitation, there is an established literature on using expert responses to inform decisions in contexts of limited data, and the particular expert-based approach used in cumulative impact mapping was evaluated in Teck et al. (29) and shown to be robust. Specifically for our study, risk criteria scores had relatively low variation across experts (standard deviation was usually less than half of the mean, and often less than a quarter of the mean for recreation, all fisheries, and all aquaculture). Additionally, experts were provided opportunities to comment and disagree with aggregated results, but in all ecosystem services, experts were satisfied with the results. Taken together (low variation across experts and no further refinement by experts), these results indicate that expert scores were relatively stable. See Table S7 for a summary of expert scores for the seven criteria across impacting activities for each ecosystem service.

### 2.4 Cumulative Impacts Model

After all ecosystem services were modeled, their spatial overlap with all activity and stressors was mapped at a 500×500m cell resolution. The spatial extent of specific ecosystem services served as the boundary for each overlapped map. All intensity data for human activities were log transformed and normalized according to the largest intensity value in each activity and stressor dataset to generate a dimensionless 0-1 intensity scale (33). Cumulative impact *I*_*c*_ was calculated for each pixel according to the established cumulative impact map formula

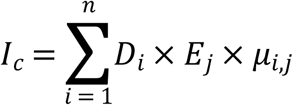

where *D*_*i*_ is the log-transformed and normalized intensity scores for activity or stressor *i*, E_j_ is the presence or absence of ecosystem service *j*, and *μ*_*i,j*_ is the risk of individual occurrences of activity or stressor *i* on ecosystem service *j* (see Supplementary Methods, 33). Cumulative impacts were calculated both including and excluding service and value dimensions to examine the contribution of service and value dimensions on ecosystem services.

### 2.5 Future Risk to Ecosystem Services

To partially assess future risks to ecosystem services, experts were asked to quantify risk to two global stressors and one regional stressor of high concern, given the changing climate and development trajectory of British Columbia. These measures of future risk were not included in the cumulative impact maps, as the maps only included risk estimates for current activities and stressors that cause impact. Experts were asked to quantify risk from sea surface temperature rise and ocean acidification according to projections for the year 2100 (3°C increase and 0.3 pH decrease, respectively, (36), and to quantify risk from a major oil spill (>40 000 m^3^, (37)). All risk scores were normalized so that the resulting expert scores were scaled between 0 – 1. These future risks were not incorporated into the cumulative impact maps.

### 2.6 Understanding Mechanisms of Impact

We asked experts in the risk survey to indicate whether or not the given activities and stressors affected their chosen ecosystem service directly or indirectly (or neither or both), with an optional follow-up to describe the pathways of impact.

## 3. Results

### 3.1 Impacts to Ecosystem Services

Our results indicated that all modeled ecosystem services are impacted across most – if not all – of their range (Figures 1 and 2). Controlling for total range, the average per-cell impact was highest for commercial demersal fisheries (*I*_*c*_ = 0.43), followed by commercial pelagic fisheries (0.41), shellfish aquaculture (0.36), finfish aquaculture (0.35), potential renewable energy (0.32), marine recreation (0.31), coastal protection (0.18), and aesthetics (0.06). Considering only ecosystem service supply dimensions, the position of potential energy and recreation are switched, otherwise this ranked list of ecosystem services facing impacts is consistent with the list considering service and value dimensions. However, all ecosystem services vary greatly in their relative impacts (Figure 3). Most ecosystem services have per-cell *I*_*c*_ values that range from ∼0-0.8, except aesthetics, which only has *I*_*c*_ values ∼0-0.4. This ranking is largely consistent with a ranked list of ecosystem services facing impact only considering ecosystem service supply dimensions.

**Figure 1.**
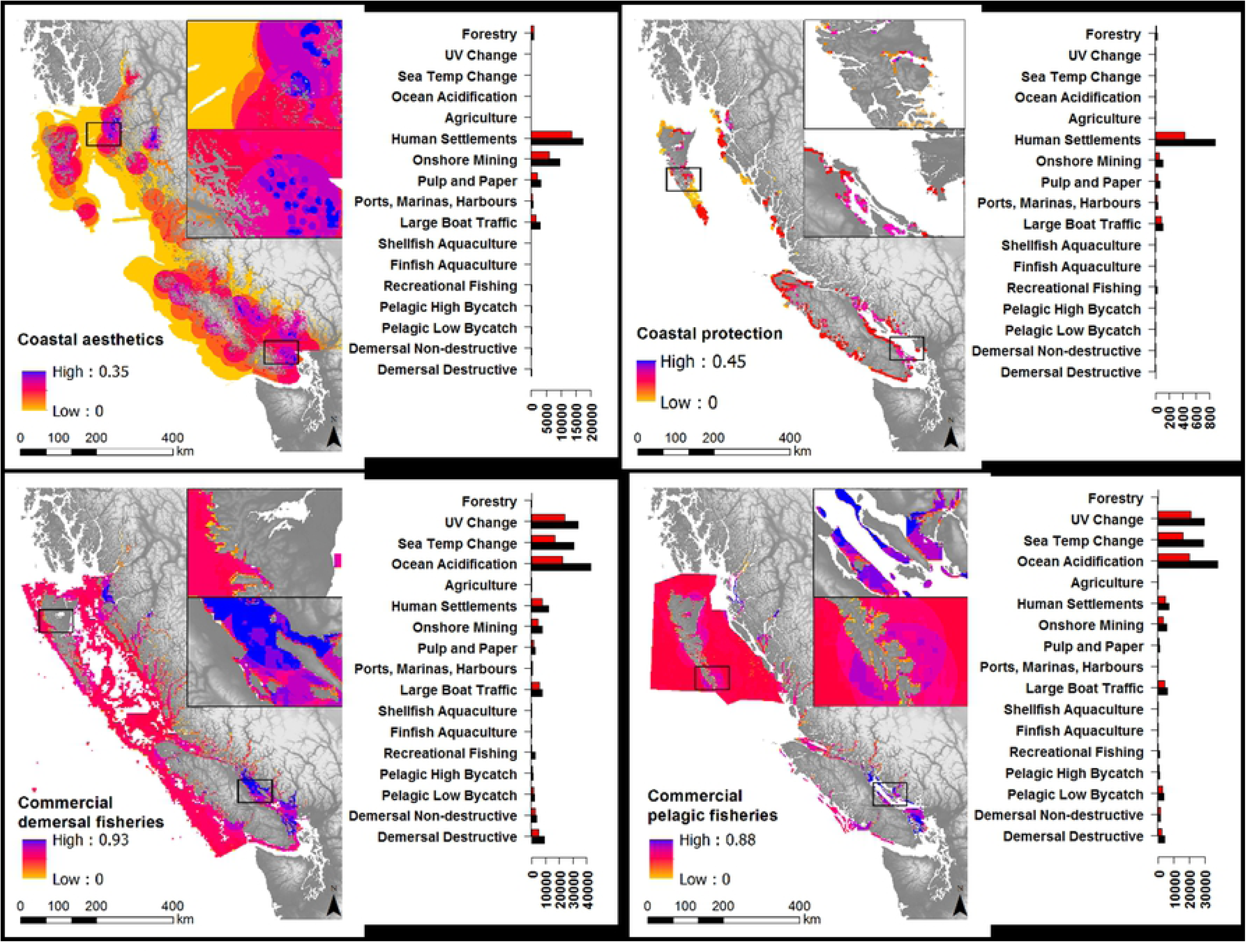
Cumulative impact maps for four ecosystem services (aesthetics, coastal protection, commercial demersal and commercial pelagic fisheries), with associated bar graphs of causes of impact. Maps display the summed impact of all drivers and stressors to each ecosystem service; bar graphs show total impact values for each activity or stressor. Red bars indicate impact only accounting for ecosystem service supply dimensions, and black bars indicate impact accounting for the entire ecosystem service cascade (supply, service, and value). Coastal protection is not to scale to allow for visibility. Four human activities and stressors that cause impact have been left off the bar graphs because they contribute negligible levels of impact across ecosystem services (small docks, log dumping, ocean dumping, and industry).

**Figure 2.**
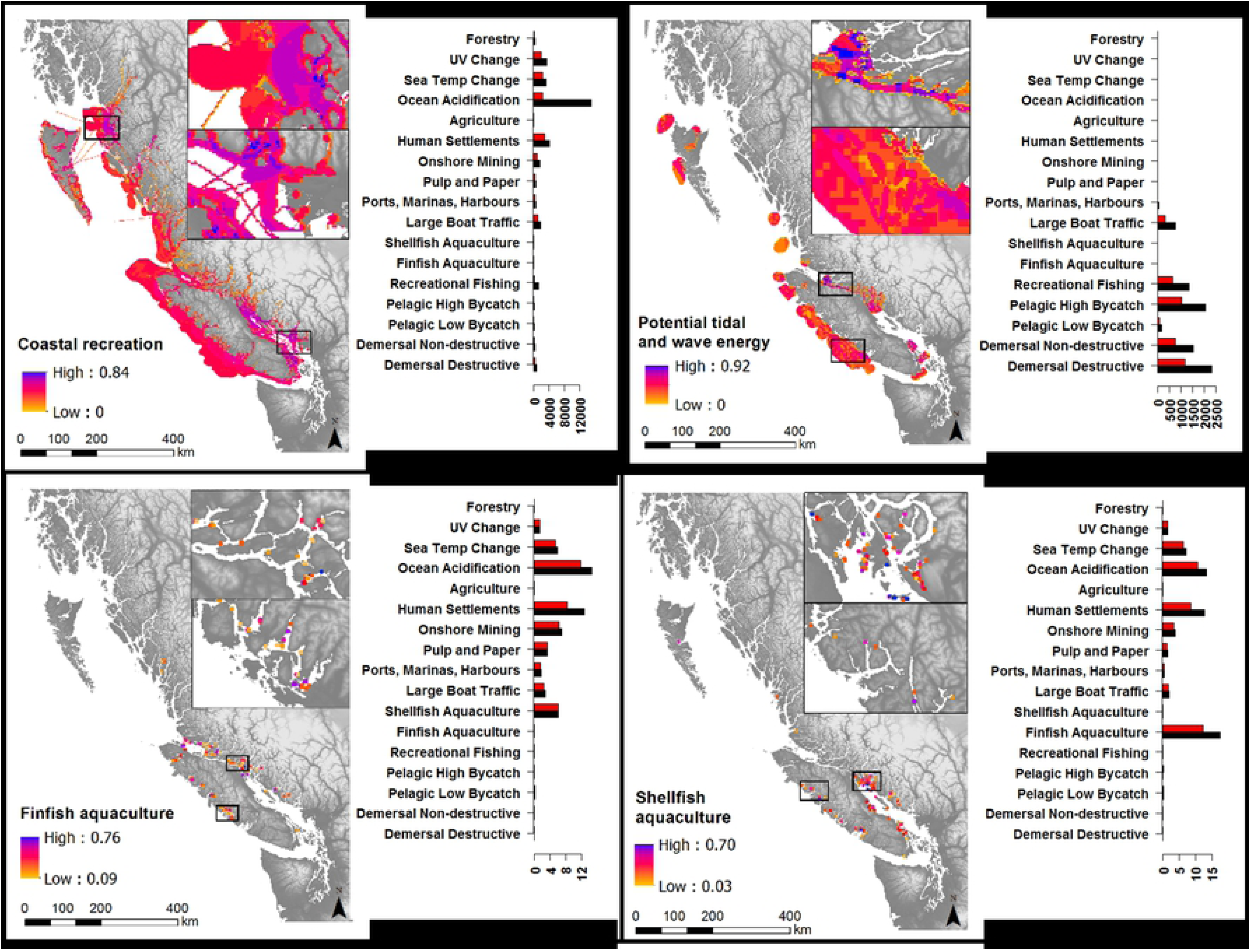
Cumulative impact maps for four ecosystem services (recreation, energy, finfish and shellfish aquaculture), with associated bar graphs of causes of impact. Maps display the summed impact of all drivers and stressors to each ecosystem service; bar graphs show total impact values for each activity or stressor. Red bars indicate impact only accounting for ecosystem service supply dimensions, and black bars indicate impact accounting for the entire ecosystem service cascade (supply, service, value). Aquaculture sites are not to scale to allow for visibility. Four human activities and stressors that cause impact have been left off the bar graphs because they contribute negligible levels of impact across ecosystem services (small docks, log dumping, ocean dumping, and industry).

**Figure 3.**
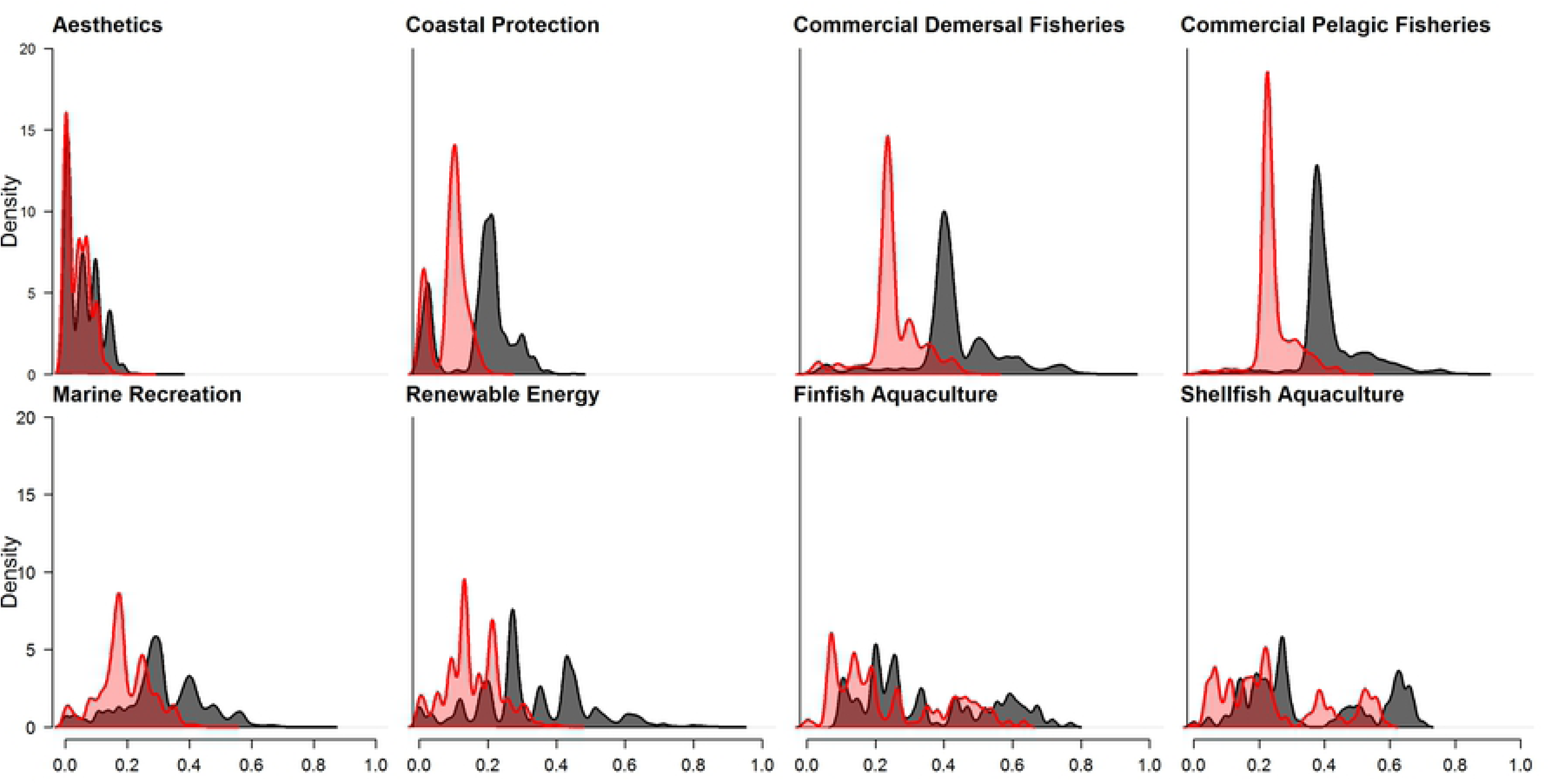
Density histograms of per-cell *I*_*c*_ values for each ecosystem service. Red histograms indicate per pixel impact only accounting for ecosystem service supply, and black histograms indicate impact accounting for all dimensions (supply, service, value).

Considering total summed impact across the range of an ecosystem service, commercial demersal fisheries face the highest impact (aggregate *I*_*c*_ = 1.57×10^5^), followed by commercial pelagic fisheries (1.27×10^5^), aesthetics (3.54×10^4^), marine recreation (3.28×10^4^), potential renewable energy generation (8.24×10^3^), coastal protection (1.24×10^3^), finfish aquaculture (59.9) and shellfish aquaculture (55.9). Considering only ecosystem service supply dimensions, the position of finfish aquaculture and shellfish aquaculture are switched; otherwise this ranked list of ecosystem services facing impacts is consistent with the ranked list when service and value dimensions are also considered. This ranking largely follows the ranking in spatial range of the ecosystem services themselves. For many ecosystem services, higher levels of impact were found on the south of the coast, between Vancouver Island and the mainland (for finfish and shellfish aquaculture and potential energy generation, and coastal protection), and the north coast (for aesthetics, coastal protection, demersal and pelagic fisheries, and marine recreation). Major hotspots of impact are similar when considering service and value dimensions of impact versus not considering them.

Different groups of drivers and activities generated prominent impacts for different ecosystem services (Figures 1 and 2). Climate related stressors contributed high levels of impact to demersal and pelagic fisheries, marine recreation, finfish aquaculture and shellfish aquaculture. Ocean acidification was the main climate related stressor contributing to impact in these ecosystem services. Climate related stressors had the highest spatial range across all ecosystem services (occupying all map cells). Land-based activities contributed high levels of impact to aesthetics, coastal protection, and both aquaculture categories. Human settlements and onshore mining contributed the most impact to most of these ecosystem services. Coastal commercial activities contributed high levels of impact to finfish aquaculture. Aquaculture was seen as a prominent activity impacting itself, as experts scored risk to ecosystem service supply, service, and value dimensions high for aquaculture, and multiple experts described the self-harmful practices and invasive and disease problems of aquaculture. They also cited the poor public attitude towards aquaculture as a high risk to itself. Fisheries contributed high levels of impact to potential tidal and wave energy. Experts scored risk to service and value dimensions from fisheries to potential energy generation high, specifically the effects of fisheries on access to good renewable energy sites.

### 3.2 Importance of Service and Value Metrics to Impact Scores

Across all ecosystem services, total and per-pixel impact scores were more severe when including risk to service and value dimensions in impact calculations than excluding them (Figures 1, 2, and 3). Resulting maps show greater overall impact across the spatial range of all ecosystem services when these service and value dimensions are included on top of ecosystem service supply dimensions (Figure S1). Though we use an additive model (and so any additional criteria will add to total impact), the service and value dimensions contributed a substantial proportion towards total impact (Figure 3). Including these service and value dimensions had the greatest proportional increase in per-pixel *I*_*c*_ for potential renewable energy generation (2.07 times greater than only considering impact on ecosystem service supply), followed by coastal protection (1.97), commercial demersal fisheries (1.70), commercial pelagic fisheries (1.69), recreation (1.64), aesthetics (1.42), finfish aquaculture (1.24), and shellfish aquaculture (1.16). Considering total *I*_*c*_ values, including service and value dimensions had the greatest proportional increase for recreation (2.50), followed by potential renewable energy generation (2.07), coastal protection (1.97), commercial demersal fisheries (1.70), commercial pelagic fisheries (1.69), aesthetics (1.42), finfish aquaculture (1.28), and shellfish aquaculture (1.19). The only case where considering impacts on service and value dimensions did not add to impact estimates was the impact of shellfish aquaculture on itself (Figure 2).

### 3.4 Future Risk

Considering future impacts, experts perceived that some ecosystem services are at greater risk from some future climate stressors than potential major oil spill, while others are at greater risk from potential major oil spills (Figure 4). Aesthetics, coastal protection, and potential energy generation were all perceived to be at higher risk from a major oil spill on the coast, and face no risk from future sea temperature or ocean acidification. Coastal protection and potential energy generation were perceived to be at high risk from sea level rise, but we did not have spatial data for this stressor so we do not represent it here. In contrast, fisheries, aquaculture, and marine recreation all appeared to be at higher risk from future ocean acidification and sea surface temperature rise, and particularly ocean acidification.

**Figure 4.**
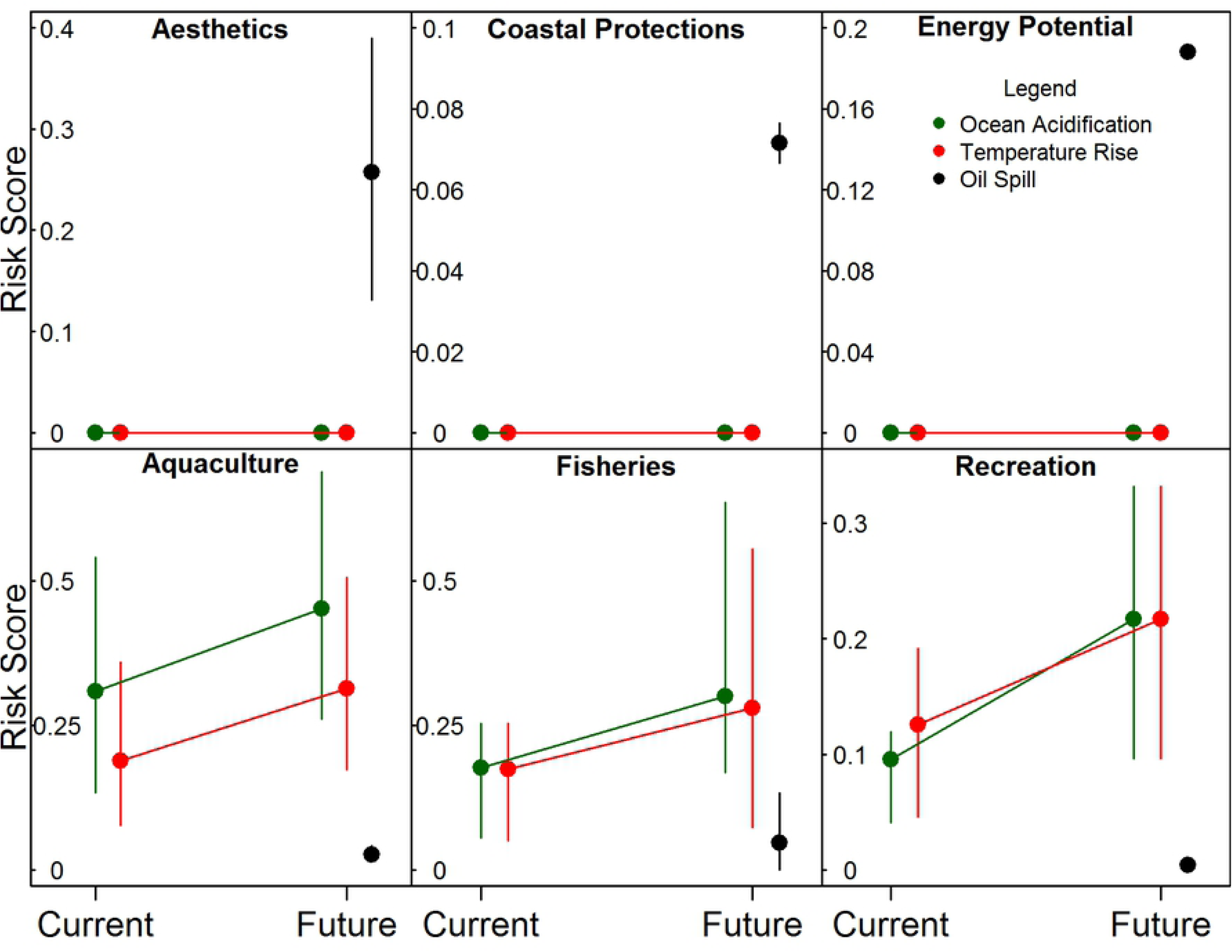
The risk posed by future climate change risks and oil spills on six ecosystem services, compared with current climate change risks. Points represent mean risk scores, error bars represent 25th and 75th percentiles, and lines connecting points demonstrate the trajectory of risk from current conditions to future conditions.

### 3.3 Relative Importance of Ecosystem Service Supply, Service, and Value for Impact

Based on expert ranking, risk to ecosystem services is dependent on diverse criteria of exposure and consequence, without a clearly dominant criteria influencing risk (Figure 5). For exposure criteria, experts considered the spatial extent of individual occurrence of activities to be most important, followed by the recovery time of an ecosystem service to an impact, and finally the frequency at which an ecosystem experiences an activity. For consequence criteria, experts considered the magnitude of change to the biophysical processes that produce the ecosystem service to be most important, followed by how the perceived quality of an ecosystem service changes in response to an impacting activity, the extent to which the environment is impacted (from individual species to entire ecosystems), and finally the changes to access to an ecosystem service. However, simple rankings mask the finding that experts perceived all criteria to contribute non-trivially to risk (the best model estimated frequency to contribute 20% to exposure, and access to contribute 19% to consequence), and that there was a diversity of weights considered across our experts (Figure 5), reflecting that some experts considered service and value dimensions of ecosystem services to be more important than biophysical supply components.

**Figure 5.**
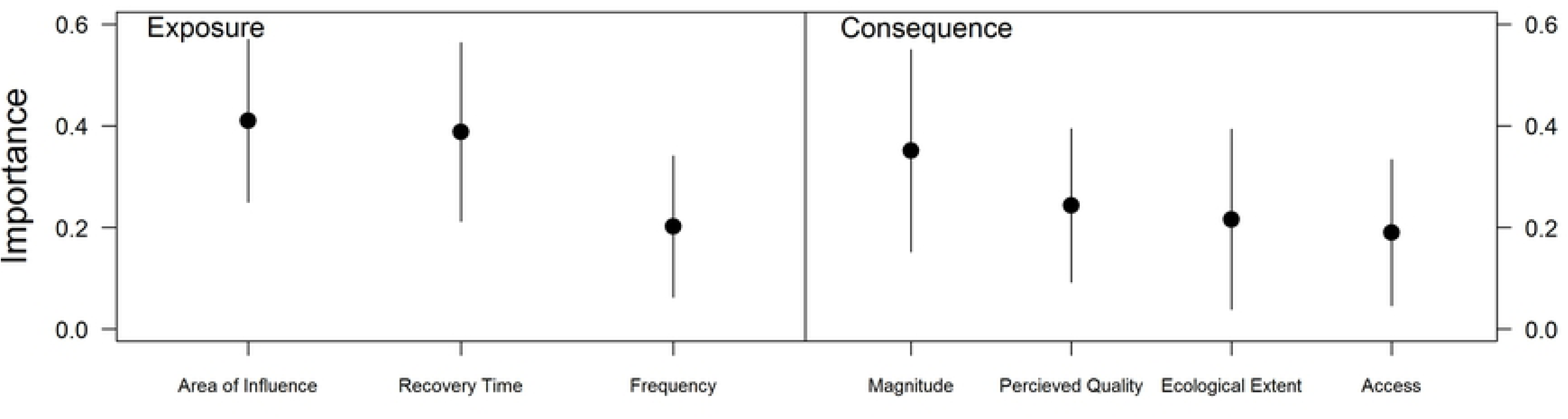
The perceived importance of risk criteria to exposure and consequence. Points and error bars represent mean and standard deviations of the distribution of relative importance of risk criteria.

### 3.4 Causal Pathways of Impacts

Experts suggested diverse prominent pathways of effect from anthropogenic activities and stressors among the ecosystem services (Figure 6). Across all types of impact, including fisheries impacts, coastal commercial activities, land based activities and climate stressors, some ecosystem services have consistent impact pathway types. Most aesthetics experts suggested that impact pathways to aesthetics are direct, with some specifically suggesting that the physical footprint of the activity is often all that matters for aesthetics. Renewable energy potential was an ecosystem service that many experts suggested was not affected by any activity or stressor, though a sizeable minority suggested that fisheries affect it directly through restricting access, and that climate change affects it both directly and indirectly through changing sea levels and affecting energy demand (which affects the infrastructural needs and suitability of locations for energy sites). Coastal protection was most often thought to be directly affected by activities through physical damage to kelp and seagrass beds and through pollution, and some suggested that recreational fishing vessels crowd estuaries and fjords, destroying habitat that support wave attenuation, and themselves generate additional wake that can risk coastlines. Most experts suggested that aquaculture is predominantly directly affected by some activities (such as land based runoff) but indirectly through others (such as invasive and disease spread from fishing vessels and ships), as well as directly and indirectly from sea temperature and ocean acidification affecting the harvested species as well as organisms that they feed on. Fisheries and recreation were both suggested to face both direct and indirect impacts from climate change, fisheries, coastal commercial, and land-based sources, according to experts. Many experts suggested that changes to foodwebs and other ecological dynamics result in indirect impacts along with direct impacts from all types of human impacts.

**Figure 6.**
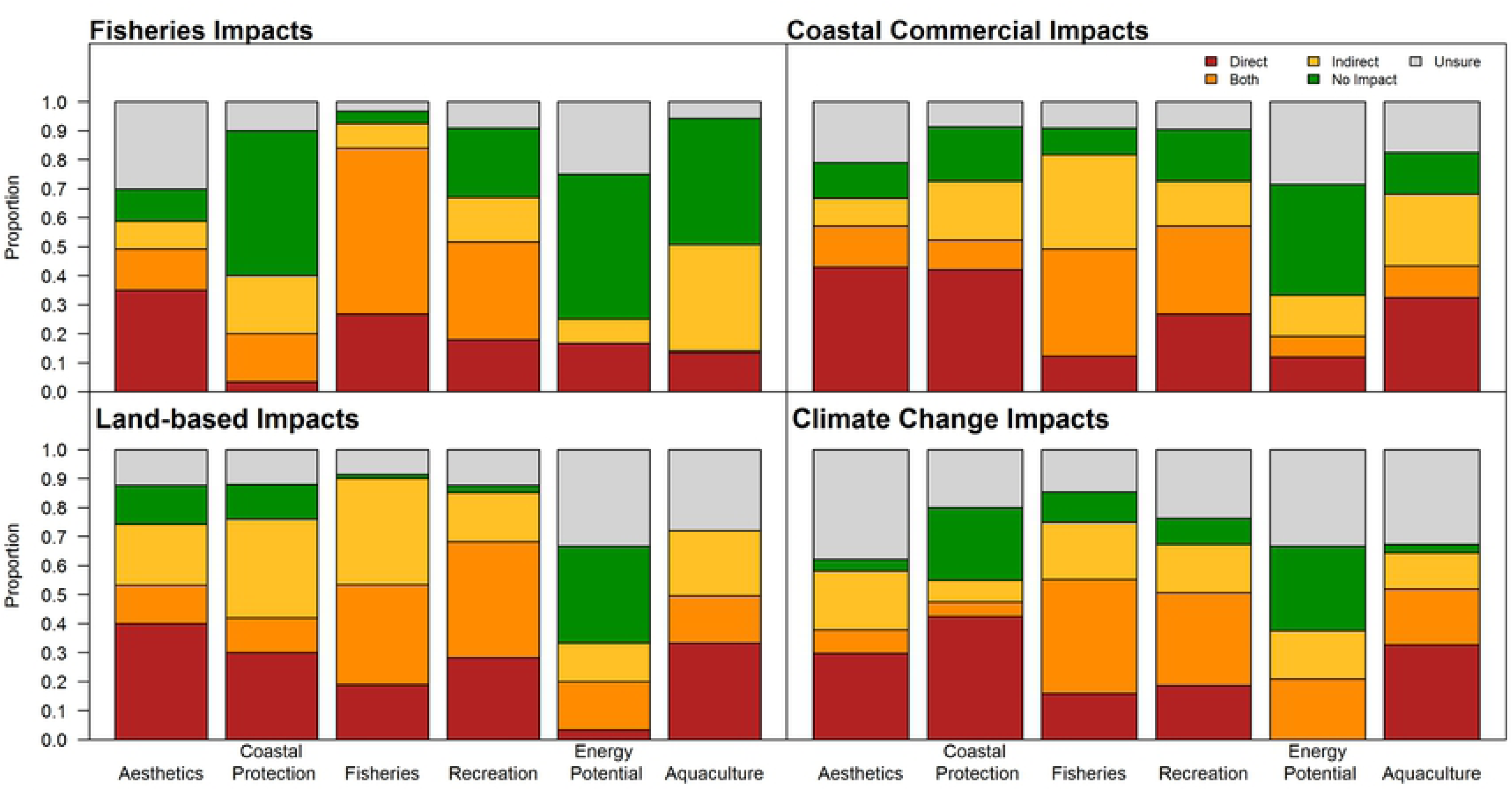
The proportion of each type of impact pathways (direct, both direct and indirect, indirect, no impact, and unsure) from four categories of activities and stressors to the eight ecosystem services, as indicated by experts.

## 4. Discussion

### 4.1 Including Service and Value Dimensions Lead to Greater Accounting of Impact

Considering service and value dimensions in addition to ecosystem service supply leads to more severe cumulative impact scores and a greater diversity of impact pathways. However, our results indicate that consideration of service and value dimensions does not greatly affect a relative understanding of impact across ecosystem services: only considering ecosystem service supply generated similar ranks or ecosystem services facing greatest impact and highlighting hotspots of impact. Our results may be interpreted to suggest that impact maps of ecosystem services that only consider supply dimensions may accurately generate conclusions about what services face greatest impact and where they face greatest impact. However, considering service and value dimensions set the scope of which services are considered for impact assessment (by determining which services are most valued) and their spatial boundaries (because people do not benefit from ecosystem services throughout their entire range of biophysical production).

Additionally, ours is an initial investigation into the importance of service and value dimensions for ecosystem service impact, and expert scores of risk criteria may fail to emphasize service and value dimensions because of two important biases. First, many of the experts taking part in our survey have ecological and biophysical training. Second, most prominent frameworks of ecosystem service change represent impacts as mediated solely through the biophysical community (19, 21, 22), which may affect how experts think about impacts. In cases where there are important impacts that overwhelmingly impact ecosystem services through service and value dimensions, excluding these dimensions may lead to different rankings of threatened ecosystem services and different map hotspots. Determining how prevalent these cases are in different settings remains to be seen. Regardless of understanding relative impact, our results indicate that studies based only on supply dimensions may underrepresent the processes that generate impact to ecosystem services. Considering the ecosystem service cascade from service supply through service delivery through satisfying values (15) may lead to a more detailed understanding about impacts and potential responses to these impacts.

### 4.2 Mapping Ecosystem Services Allows for Insights Not Afforded by Mapping Habitats

Impact on ecosystem services is a function not only of spatial overlap with concurrent activities and stressors, but of the risk through ecosystem service supply, service, and value dimensions of ecosystem services as well. Many ecosystem services face high impact in the area between Vancouver Island and the mainland, a finding reinforced by previous studies that focused solely on impacts to habitats (4, 8, 10, 28, 38). Not all ecosystem services have impact hotspots here, however, reflecting the importance of accurately mapping ecosystem services. While our study alongside previous ones may share similar patterns of human activity, the distribution of ecosystem services themselves is important in determining where areas of high impact are. The marine InVEST models use data of environmental process and human activity to spatially represent ecosystem services, allowing us to directly model ecosystem services (17).

Accurately representing the overlap of activities and stressors on ecosystem services generate additional insights. Knowing where ecosystem services are at highest risk can allow managers to assess impact relative to areas of high demand (24). Areas of high risk to coastal protection were concentrated close to population centers (in the southern Strait of Georgia), partly because the human activities that might benefit the most from coastal protection – human settlements – also provide the largest impact to coastal protection. Spatial representation also allows for an understanding of whether an ecosystem service faces high risk on account of large spatial range despite low per-area impact (such as aesthetics), versus ecosystem services that face low total impact because of limited geographic range despite having high per area impact (such as shellfish and finfish aquaculture). Aesthetics was found to be the least impacted ecosystem service per unit area, indicating that a beautiful coast may mask a highly impacted coast.

Explicit inclusion of risk criteria (exposure and consequence) is important because activities and stressors with extensive spatial range and high overlap with ecosystem services do not necessarily generate high impact. Similar to a recent cumulative impact mapping study on coastal ecosystems in British Columbia (10), and along the California current (4) we found climate change impacts to be important stressors (especially ocean acidification), highlighting the importance of their inclusion in analysis, and cautioning results from mapping studies that do not include them (8). For example, ecosystem services dependent on invertebrate and finfish (fisheries, aquaculture, and marine recreation) were highly impacted by ocean acidification. Climate change stressors exist across the entire marine system along the British Columbia coast and consequently fully overlap with every ecosystem service we modeled, yet impact on some ecosystem services is driven largely by climate impacts (such as fisheries and aquaculture) while others are largely indifferent to climate change stressors (such as aesthetics, coastal protection and potential energy generation). Indeed, kelp and seagrasses associated with coastal protection may benefit with ocean acidification (39, 40). Sea-level rise was indicated as a high risk stressor especially to coastal protection, but we did not have spatial data for sea level rise. While the risk scores for potential energy generation are uncertain given the input from only one expert, the conclusion that energy generating infrastructure and planning faces greater risk from human activities than climate impacts is plausible.

This dichotomy between global and regional impacts may exacerbate in the future, as experts suggested that future climate change impacts (specifically warming and ocean acidification) will be a higher risk to those climate-sensitive ecosystem services compared to current conditions, while climate-insensitive ecosystem services will face similar risk levels. For these latter ecosystem services, potential future development may pose a greater cause for concern. Future oil spill potential related to planned developments of oil and gas with associated marine shipping poses a significant risk to these ecosystem services. Previous efforts to compare climate change impacts with future developments in British Columbia indicated that climate change has greater regional scale impact across ecosystem types but lower local impact (10). We show that some ecosystem services – in contrast to ecosystem types – show varying degrees of risk to different types of stressors, leading to insensitivity to climate change stressors for some ecosystem services at local and regional scales.

### 4.3 Service and Value Dimensions Are Important for Understanding Causes of Impact

Experts in our survey treated individual service and value dimensions with comparable importance to supply dimensions when ranking scenarios. Service and value dimensions are definitional to ecosystem services yet are often overlooked in quantitative assessments. Despite supply dimensions potentially being sufficient for understanding which ecosystem services are most impacted relative to one another, relying on supply dimensions alone is shortsighted for two reasons. First, any quantitative measure of impact is likely to be an underestimate (13). Service and value factors, such as how people perceive an ecosystem service, can regulate the extent to which people enjoy and benefit from the ecosystem service (23). For example, open-pen finfish aquaculture practices are perceived negatively by many people in British Columbia (26), creating a self-stigmatized industry. Whether public perceptions on finfish aquaculture are warranted or not, they affect aquaculture as the aquaculture industry has launched marketing campaigns to fight its reputation (www.bcsalmonfacts.ca). Second, many ecosystem services can be impacted largely (even solely) through changes to access and perceived value. Experts indicated that potential wave and tidal energy production face risk from fisheries and ports partly through the competition for space, as access to suitable power generation sites can be blocked or zoned out by competing interests for the area. If situations where impacts occur through service and value dimensions become more common then relying on supply dimensions may no longer be suitable for understanding which ecosystem services face highest relative impact.

### 4.4 Accounting for Pathways of Impact Can Improve Cumulative Impact Models

Ecosystem services may require different data and representation techniques than ecosystem types. Unlike ecosystem types, ecosystem services are not variants of geographical classes; ecosystem services do not only exist on a landscape but are related to people’s values and ability to obtain them (23, 41). The same activity may have different impact pathways on two different ecosystem services because one ecosystem service could be primarily impacted through a change in species density while another could be impacted primarily because the activity restricts people to a region through property rights and trespassing laws. The greater diversity of potential pathways of impact that ecosystem services face arguably puts greater emphasis on understanding the causal processes of impact for ecosystem services than for ecosystem types.

Given the diverse kinds of ecosystem services that exist, a common spatial representation of specific human activities and stressors across ecosystem services may produce misleading results in two important ways. First, the impact pathway important for the ecosystem service should dictate the size of the zone of influence (8). Many experts in our study suggested that aesthetics are directly impacted from most activities and stressors, and that what matters is the physical footprint of any activity. We have onshore mining spatially represented to account for acid mine drainage and tailings that occur kilometers away from mines themselves. This area of influence is likely appropriate when mapping impact to ecosystem services affected by these processes, such as fisheries and aquaculture, but it may lead to overestimated overlap of mining impacts and aesthetics. Future efforts to map impacts on ecosystem services should match the spatial representation of activities with relevant impacts. Second, not all experts understand the impact pathways the same, which means they may not answer the same questions. The precedent set here using expert surveys, in conjunction with impact mapping, asks experts to assess vulnerability/risk to an activity “considering all relevant impacts”. This open question framing allows for a tractable survey, yet our results suggest that what is considered in “all relevant impacts” may vary from expert to expert for a given human activity. What’s hidden in our resulting maps is a significant epistemic uncertainty that can be reduced with appropriate elicitation strategies (42). Future expert elicitation processes should emphasize specific pathways when assessing risk, even if it means batching surveys into sets of different impact pathways so different experts quantify risk to different impacts.

### 4.5 Limitations and Opportunities

While we present advancement in cumulative impact mapping – namely representing ecosystem services and accounting for impact along supply, service, and value dimensions – and recommend data considerations specifically for ecosystem services, we must also acknowledge persistent limitations of impact mapping. Most importantly these include a static representation of impact and a simplistic model of cumulative impacts (3, 8, 28). Though experts considered temporal criteria of exposure as less important than area of influence, they were still important components of risk, showing that temporal considerations are essential. Spatial models are often snapshots in time, and though we include some temporal dynamics (assessing risk of foreseen impacts) there are many important temporal aspects of impact that are not captured. We do not represent future impacts spatially (but see Murray et al. (10) for a spatial analysis of proposed projects), though understanding future impacts would be highly valuable to managers. We also do not account for historic impacts. By focusing on contemporary impacts we set a contemporary benchmark and do not consider change from ecosystem service states that may be more ideal, such as times in the past when overfishing was not as prevalent (43, 44).

The cumulative impact model we employ assumes an additive, relative model of impact with no upper bound. Both activity intensity and risk scores were normalized between 0-1, so components of the model have measurement boundaries, but the cumulative impact can aggregate indefinitely. Empirical studies have shown additive cumulative impacts to occur in a minority of situations (45, 46). Synergistic impacts – when the total impact is greater than the sum of component impacts – occur often, especially when more than two impacts co-occur (5, 45, 47). Antagonistic impact – when total impact is less than the sum of component impacts – are also prevalent, and have been shown when global impacts interact with local impacts (48, 49). The theoretically limitless measure of impact produced by the model employed here also assumes that impacts can accumulate indefinitely, and that thresholds do not exist (3). These are obviously false assumptions, but this model can still provide broad insights into the relative impact faced by multiple ecosystem services.

Finally, this work depends on input from experts. Expert input can be affected by biases (34) and therefore can increase uncertainty of results. Uncertainty in expert responses is even higher when few experts provide input (such as for potential wave and tidal energy generation here), and responses should be considered as hypotheses requiring empirical validation (6). However, where empirical data does not exist (such as in the case here), expert input elicited through structured processes can provide valuable input for decision-making (25, 30, 34), including the specific elicitation techniques used here (29).

Despite modeling limitations, mapping cumulative impacts to ecosystem services allows for unique planning opportunities. Ocean managers can use this approach to explore the spatial feasibility of potential coastal uses, as we show for potential wave and tidal power generation, even if precise risk estimates are not available. By mapping areas of potential energy generation, we see that the areas of lowest threat to energy generation are the central coast and some areas between Vancouver Island and the mainland. These are relatively unpopulated areas, which may mean higher infrastructure costs to establish turbines, but these costs may be worth avoiding impediments in more populous areas.

## 5. Conclusion

By mapping cumulative impacts to ecosystem services, we can better steward our ecosystems and understand the dual relationship of humans to the environment: as agents of change and beneficiaries of services (12). We have demonstrated the kinds of rich insights that can be gained from mapping impacts to ecosystem services, including: 1) discovering where, and by what means, different ecosystem services face the greatest impact; 2) determining what ecosystem services are comparatively worse (or better) off under current conditions; 3) understanding the ways in which impacts manifest; 4) assessing spatial feasibility for new ocean uses. We have also demonstrated the importance of considering service and value dimensions in assessing impact. We argue that considering service and value dimensions is not only important to more fully understand impact, but also to plan effective management responses. Finally, we have pointed to areas of future methodological refinement, and encourage greater innovation in cumulative impact mapping. Ecosystem services can be highly location specific (23), so future risk assessments are warranted in new places. Understanding risk and impact to ecosystem services should be an essential management priority to maintain the flow of services we benefit from.

## Acknowledgements

We are grateful to A. Thompson and T. Coyle for expert identification and help building the online survey. We would also like to thank G. Peterson, M. O’Connor, and S. Gergel for reviewing and commenting on the work. We would also like to thank WWF Canada for the use of data on human activities. GGS benefitted from NSERC’s Michael Smith Foreign Study Supplement and support from the Pacific Institute for Climate Solutions to facilitate this work. This work was also supported by the David and Lucile Packard Foundation.

## Literature Cited

1. Arkema KK, Abramson SC, Dewsbury BM. Marine ecosystem-based management: from characterization to implementation. Frontiers in Ecology and the Environment. 2006;4(10):525–32.

2. Granek EF, Polasky S, Kappel CV, Reed DJ, Stoms DM, Koch EW, et al. Ecosystem Services as a Common Language for Coastal Ecosystem-Based Management. Conservation Biology. 2009;24(1):207–16.

3. Halpern BS, Fujita R. Assumptions, challenges, and future directions in cumulative impact analysis. Ecosphere. 2013 2013/10/01;4(10):art131.

4. Halpern BS, Kappel CV, Selkoe KA, Micheli F, Ebert CM, Kontgis C, et al. Mapping cumulative human impacts to California Current marine ecosystems. Conservation letters. 2009;2(3):138–48.

5. Halpern BS, McLeod KL, Rosenberg AA, Crowder LB. Managing for cumulative impacts in ecosystem-based management through ocean zoning. Ocean & Coastal Management. 2008;51(3):203–11.

6. Micheli F, Halpern BS, Walbridge S, Ciriaco S, Ferretti F, Fraschetti S, et al. Cumulative human impacts on mediterranean and black sea marine ecosystems: assessing current pressures and opportunities. PLoS ONE. 2013;8(12):e79889.

7. Selkoe KA, Halpern BS, Ebert CM, Franklin EC, Selig ER, Casey KS, et al. A map of human impacts to a “pristine” coral reef ecosystem, the Papahānaumokuākea Marine National Monument. Coral Reefs. 2009 2009/09/01;28(3):635–50.

8. Ban NC, Alidina HM, Ardron JA. Cumulative impact mapping: Advances, relevance and limitations to marine management and conservation, using Canada’s Pacific waters as a case study. Marine Policy. 2010;34(5):876–86.

9. Clark D, Goodwin E, Sinner J, Ellis J, Singh G. Validation and limitations of a cumulative impact model for an estuary. Ocean & Coastal Management. 2016;120:88–98.

10. 10. Murray CC, Agbayani S, Ban NC. Cumulative effects of planned industrial development and climate change on marine ecosystems. Global Ecology and Conservation. 2015;4:110–6.

11. Allan JD, McIntyre PB, Smith SDP, Halpern BS, Boyer GL, Buchsbaum A, et al. Joint analysis of stressors and ecosystem services to enhance restoration effectiveness. Proceedings of the National Academy of Sciences. 2013 January 2, 2013;110(1):372–7.

12. MA. Ecosystems and human well-being: Island press Washington, DC:; 2005.

13. Tallis H, Lester SE, Ruckelshaus M, Plummer M, McLeod K, Guerry A, et al. New metrics for managing and sustaining the ocean’s bounty. Marine Policy. 2012;36(1):303–6.

14. Oliveira J, Cunha A, Castilho F, Romalde J, Pereira M. Microbial contamination and purification of bivalve shellfish: Crucial aspects in monitoring and future perspectives–A minireview. Food Control. 2011;22(6):805–16.

15. Haines-Young R, Potschin M. The links between biodiversity, ecosystem services and human well-being. Ecosystem Ecology: a new synthesis. 2010:110–39.

16. Angradi TR, Launspach JJ, Bolgrien DW, Bellinger BJ, Starry MA, Hoffman JC, et al. Mapping ecosystem service indicators in a Great Lakes estuarine Area of Concern. Journal of Great Lakes Research. 2016.

17. Guerry AD, Ruckelshaus MH, Arkema KK, Bernhardt JR, Guannel G, Kim CK, et al. Modeling benefits from nature: using ecosystem services to inform coastal and marine spatial planning. International Journal of Biodiversity Science, Ecosystem Services & Management. 2012;8(1-2):107–21.

18. Sharp R, Tallis H, Ricketts T, Guerry A, Wood S, Chaplin-Kramer R, et al. InVEST user’s guide. The Natural Capital Project, Stanford. 2014.

19. Rounsevell MDA, Dawson T, Harrison P. A conceptual framework to assess the effects of environmental change on ecosystem services. Biodiversity and Conservation. 2010;19(10):2823–42.

20. Mach ME, Martone RG, Chan KM. Human impacts and ecosystem services: Insufficient research for trade-off evaluation. Ecosystem Services. 2015;16:112–20.

21. Collins SL, Carpenter SR, Swinton SM, Orenstein DE, Childers DL, Gragson TL, et al. An integrated conceptual framework for long-term social-ecological research. Frontiers in Ecology and the Environment. 2010;9(6):351–7.

22. Kelble CR, Loomis DK, Lovelace S, Nuttle WK, Ortner PB, Fletcher P, et al. The EBM-DPSER Conceptual Model: Integrating Ecosystem Services into the DPSIR Framework. PLoS ONE. 2013;8(8):e70766.

23. Chan KM, Guerry AD, Balvanera P, Klain S, Satterfield T, Basurto X, et al. Where are cultural and social in ecosystem services? A framework for constructive engagement. BioScience. 2012;62(8):744–56.

24. Wieland R, Ravensbergen S, Gregr EJ, Satterfield T, Chan KM. Debunking trickle-down ecosystem services: The fallacy of omnipotent, homogeneous beneficiaries. Ecological Economics. 2016;121:175–80.

25. Singh GG, Sinner J, Ellis J, Kandlikar M, Halpern BS, Satterfield T, et al. Group elicitations yield more consistent, yet more uncertain experts in understanding risks to ecosystem services in New Zealand bays. PloS one. 2017;12(8):e0182233.

26. Klain SC, Chan KMA. Navigating coastal values: Participatory mapping of ecosystem services for spatial planning. Ecological Economics. 2012;82(0):104–13.

27. Dawson M. Selling British Columbia: tourism and consumer culture, 1890-1970: UBC Press; 2005.

28. Murray CC, Agbayani S, Alidina HM, Ban NC. Advancing marine cumulative effects mapping: An update in Canada’s Pacific waters. Marine Policy. 2015;58:71–7.

29. Teck SJ, Halpern BS, Kappel CV, Micheli F, Selkoe KA, Crain CM, et al. Using expert judgment to estimate marine ecosystem vulnerability in the California Current. Ecological Applications. 2010;20(5):1402–16.

30. Singh GG, Sinner J, Ellis J, Kandlikar M, Halpern BS, Satterfield T, et al. Mechanisms and risk of cumulative impacts to coastal ecosystem services: An expert elicitation approach. Journal of environmental management. 2017;199:229–41.

31. BCMCA. BC Marine Conservation Analysis. 2016 [cited 2016 2016]; Available from: http://bcmca.ca/.

32. GeoBC. GeoBC. 2016 [cited 2016 2016]; Available from: http://geobc.gov.bc.ca/.

33. Halpern BS, Walbridge S, Selkoe KA, Kappel CV, Micheli F, D’Agrosa C, et al. A Global Map of Human Impact on Marine Ecosystems. Science. 2008 February 15, 2008;319(5865):948–52.

34. McBride MF, Burgman MA. What Is Expert Knowledge, How Is Such Knowledge Gathered, and How Do We Use It to Address Questions in Landscape Ecology? Expert Knowledge and its Application in Landscape Ecology. 2012:11–38.

35. Halpern BS, Selkoe KA, Micheli F, Kappel CV. Evaluating and ranking the vulnerability of global marine ecosystems to anthropogenic threats. Conservation Biology. 2007;21(5):1301–15.

36. Stocker TF. Climate change 2013: the physical science basis: Working Group I contribution to the Fifth assessment report of the Intergovernmental Panel on Climate Change: Cambridge University Press; 2014.

37. Peterson CH, Rice SD, Short JW, Esler D, Bodkin JL, Ballachey BE, et al. Long-term ecosystem response to the Exxon Valdez oil spill. Science. 2003;302(5653):2082–6.

38. Ban N, Alder J. How wild is the ocean? Assessing the intensity of anthropogenic marine activities in British Columbia, Canada. Aquatic Conservation: Marine and Freshwater Ecosystems. 2008;18(1):55–85.

39. Kroeker KJ, Kordas RL, Crim R, Hendriks IE, Ramajo L, Singh GS, et al. Impacts of ocean acidification on marine organisms: quantifying sensitivities and interaction with warming. Global Change Biology. 2013;19(6):1884–96.

40. Kroeker KJ, Kordas RL, Crim RN, Singh GG. Meta-analysis reveals negative yet variable effects of ocean acidification on marine organisms. Ecology letters. 2010;13(11):1419–34.

41. Chan KM, Satterfield T, Goldstein J. Rethinking ecosystem services to better address and navigate cultural values. Ecological Economics. 2012;74:8–18.

42. Regan HM, Colyvan M, Burgman MA. A taxonomy and treatment of uncertainty for ecology and conservation biology. Ecological Applications. 2002;12(2):618–28.

43. Pauly D. Anecdotes and the shifting baseline syndrome of fisheries. Trends in ecology & evolution. 1995;10(10):430.

44. Pinnegar JK, Engelhard GH. The ‘shifting baseline’phenomenon: a global perspective. Reviews in Fish Biology and Fisheries. 2008;18(1):1–16.

45. Crain CM, Kroeker K, Halpern BS. Interactive and cumulative effects of multiple human stressors in marine systems. Wiley-Blackwell; 2008. p. 1304–15.

46. Darling ES, Côté IM. Quantifying the evidence for ecological synergies. Ecology Letters. 2008;11(12):1278–86.

47. Harley CDG, Randall Hughes A, Hultgren KM, Miner BG, Sorte CJB, Thornber CS, et al. The impacts of climate change in coastal marine systems. Ecology letters. 2006;9(2):228–41.

48. Brown CJ, Saunders MI, Possingham HP, Richardson AJ. Interactions between global and local stressors of ecosystems determine management effectiveness in cumulative impact mapping. Diversity and Distributions. 2014;20(5):538–46.

49. Brown CJ, Saunders MI, Possingham HP, Richardson AJ. Managing for Interactions between Local and Global Stressors of Ecosystems. PLoS ONE. 2013;8(6):e65765.

